# Inference of Cell Type Composition from Human Brain Transcriptomic Datasets Illuminates the Effects of Age, Manner of Death, Dissection, and Psychiatric Diagnosis

**DOI:** 10.1101/089391

**Authors:** Megan Hastings Hagenauer, Anton Schulmann, Jun Z. Li, Marquis P. Vawter, David M. Walsh, Robert C. Thompson, Cortney A. Turner, William E. Bunney, Richard M. Myers, Jack D. Barchas, Alan F. Schatzberg, Stanley J. Watson, Huda Akil

## Abstract

Psychiatric illness is unlikely to arise from pathology occurring uniformly across all cell types in affected brain regions. Despite this, transcriptomic analyses of the human brain have typically been conducted using macro-dissected tissue due to the difficulty of performing single-cell type analyses with donated post-mortem brains. To address this issue statistically, we compiled a database of several thousand transcripts that were specifically-enriched in one of 10 primary cortical cell types in previous publications. Using this database, we predicted the relative cell type composition for 833 human cortical samples using microarray or RNA-Seq data from the Pritzker Consortium (GSE92538) or publicly-available databases (GSE53987, GSE21935, GSE21138, CommonMind Consortium). These predictions were generated by averaging normalized expression levels across transcripts specific to each cell type using our R-package *BrainInABlender* (validated and publicly-released: https://github.com/hagenaue/BrainInABlender). Using this method, we found that the principal components of variation in the datasets strongly correlated with the neuron to glia ratio of the samples.

This variability was not simply due to dissection – the relative balance of brain cell types appeared to be influenced by a variety of demographic, pre- and post-mortem variables. Prolonged hypoxia around the time of death predicted increased astrocytic and endothelial gene expression, illustrating vascular upregulation. Aging was associated with decreased neuronal gene expression. Red blood cell gene expression was reduced in individuals who died following systemic blood loss. Subjects with Major Depressive Disorder had decreased astrocytic gene expression, mirroring previous morphometric observations. Subjects with Schizophrenia had reduced red blood cell gene expression, resembling the hypofrontality detected in fMRI experiments. Finally, in datasets containing samples with especially variable cell content, we found that controlling for predicted sample cell content while evaluating differential expression improved the detection of previously-identified psychiatric effects. We conclude that accounting for cell type can greatly improve the interpretability of transcriptomic data.

## 1. Introduction

The human brain is a remarkable mosaic of diverse cell types stratified into rolling cortical layers, arching white matter highways, and interlocking deep nuclei. In the past decade, we have come to recognize the importance of this cellular diversity in even the most basic neural circuits. At the same time, we have developed the capability to comprehensively measure the thousands of molecules essential for cell function. These insights have provided conflicting priorities within the study of psychiatric illness: do we carefully examine individual molecules within their cellular and anatomical context or do we extract transcript or protein en masse to perform large-scale unbiased transcriptomic or proteomic analyses? In rodent models, researchers have escaped this dilemma by a boon of new technology: single cell laser capture, cell culture, and cell-sorting techniques can provide sufficient extract for transcriptomic and proteomic analyses. However, single cell type analyses of the human brain are far more challenging (1–3) – live tissue is only available in the rarest of circumstances and intact single cells are difficult to dissociate from post-mortem tissue without intensive procedures like laser capture microscopy.

Therefore, to date, the vast majority of unbiased transcriptomic analyses of the human brain have been conducted using macro-dissected, cell-type heterogeneous tissue. On Gene Expression Omnibus (GEO) alone, there are at least 63* publicly-available macro-dissected post-mortem human brain tissue datasets, and many others are available to researchers via privately-funded portals (Stanley Medical Research Institute, Allen Brain Atlas, CommonMind Consortium). These datasets have provided us with novel hypotheses (e.g., (4,5)), but often a relatively small number of candidate molecules survive analysis despite careful sample collection, and interpreting molecular results in isolation from their respective cellular context can be exceedingly difficult. At the core of this issue is the inability to differentiate between (1) alterations in gene expression that reflect an overall disturbance in the relative ratio of the different cell types comprising the tissue sample, and (2) intrinsic dysregulation of one or more cell types, indicating perturbed biological function.

In this manuscript, we present results from an easily accessible solution to this problem that allows researchers to statistically estimate the relative number or transcriptional activity of particular cell types in macro-dissected human brain transcriptomic data by tracking the collective rise and fall of previously identified cell type specific transcripts. Similar techniques have been used to successfully predict cell type content in human blood samples (6–9), as well as diseased and aged brain samples (10–12). Our method was specifically designed for application to large, highly-normalized human brain transcriptional profiling datasets, such as those commonly used by neuroscientific research bodies such as the Pritzker Neuropsychiatric Research Consortium and the Allen Brain Institute.

We took advantage of a series of newly available data sources depicting the transcriptome of known cell types, and applied them to infer the relative balance of cell types in our tissue samples. We draw from seven large studies detailing cell-type specific gene expression in a wide variety of cells in the forebrain and cortex (2,13–18). Our analyses include all major categories of cortical cell types (17), including two overarching categories of neurons that have been implicated in psychiatric illness (19): projection neurons, which are large, pyramidal, and predominantly excitatory, and interneurons, which are small and predominantly inhibitory (20). These are accompanied by three prevalent forms of glia that make up the majority of cells in the brain: oligodendrocytes, which provide the insulating myelin sheath for axons (21), astrocytes, which help create the blood-brain barrier and provide structural and metabolic support for neurons (21), and microglia, which serve as the brain’s resident macrophages and provide an active immune response (21). We also incorporate vascular cell types: endothelial cells, which line the interior surface of blood vessels, and mural cells (smooth muscle cells and pericytes), which regulate blood flow (22). We included progenitor cells because they are widely implicated in the pathogenesis of mood disorders (23). Within the cortex, these cells mostly take the form of immature oligodendrocytes (17). Finally, the primary cells found in blood, erythrocytes or red blood cells (RBCs), carry essential oxygen throughout the brain. These cells lack a cell nucleus and do not generate new RNA, but still contain an existing, highly-specialized transcriptome (24). The relative presence of these cells could arguably represent overall blood flow, the functional marker of regional neural activity traditionally used in human imaging studies.

To characterize the balance of these cell types in psychiatric samples, we first demonstrate that our method of summarizing cell type specific gene expression into a single metric (“cell type index”) can reliably predict relative cell type balance in a variety of validation datasets. Then we discover that the predicted cell type balance of samples can explain a large percentage of the variation in macro-dissected human brain microarray and RNA-Seq datasets. This variability is driven by pre- and post-mortem subject variables, such as age, aerobic environment, and large scale blood loss, in addition to dissection. Finally, we demonstrate that our method enhances our ability to discover and interpret psychiatric effects in human transcriptomic datasets, uncovering previously-documented changes in cell type balance in relationship to Major Depressive Disorder and Schizophrenia and potentially increasing our sensitivity to detect genes with previously-identified relationships to Bipolar Disorder and Schizophrenia in datasets that contain samples with highly-variable cell content.

## 2. Methods

### 2.1 Compiling a Database of Cell Type Specific Transcripts

To perform this analysis, we compiled a database of several thousand transcripts that were specifically-enriched in one of nine primary brain cell types within seven published single-cell or purified cell type transcriptomic experiments using mammalian brain tissues (2,13–18). These primary brain cell types included six types of support cells: astrocytes, endothelial cells, mural cells, microglia, immature and mature oligodendrocytes, as well as two broad categories of neurons (interneurons and projection neurons). We also included a category for neurons that were extracted without purification by subtype (“neuron_all”). The experimental and statistical methods for determining whether a transcript was enriched in a particular cell type varied by publication (**Table 1**), and included both RNA-Seq and microarray datasets. We focused on cell-type specific transcripts identified using cortical or forebrain samples because the data available for these brain regions was more plentiful than for the deep nuclei or the cerebellum. In addition, we artificially generated a list of 17 transcripts specific to erythrocytes by searching Gene Card for erythrocyte and hemoglobin-related genes (http://www.genecards.org/).

In all, we curated gene expression signatures for 10 cell types expected to account for most of the cells in the cortex. Our final database included 2499 unique human-derived or orthologous (as predicted by HCOP using 11 available databases: http://www.genenames.org/cgi-bin/hcop) transcripts, with a focus on coding varieties. We have made this database publicly available within **Suppl. Table 1**. An updateable version is also accessible within our R package (https://github.com/hagenaue/BrainInABlender) and as a downloadable spreadsheet (https://sites.google.com/a/umich.edu/megan-hastings-hagenauer/home/cell-type-analysis).

**Table 1.**
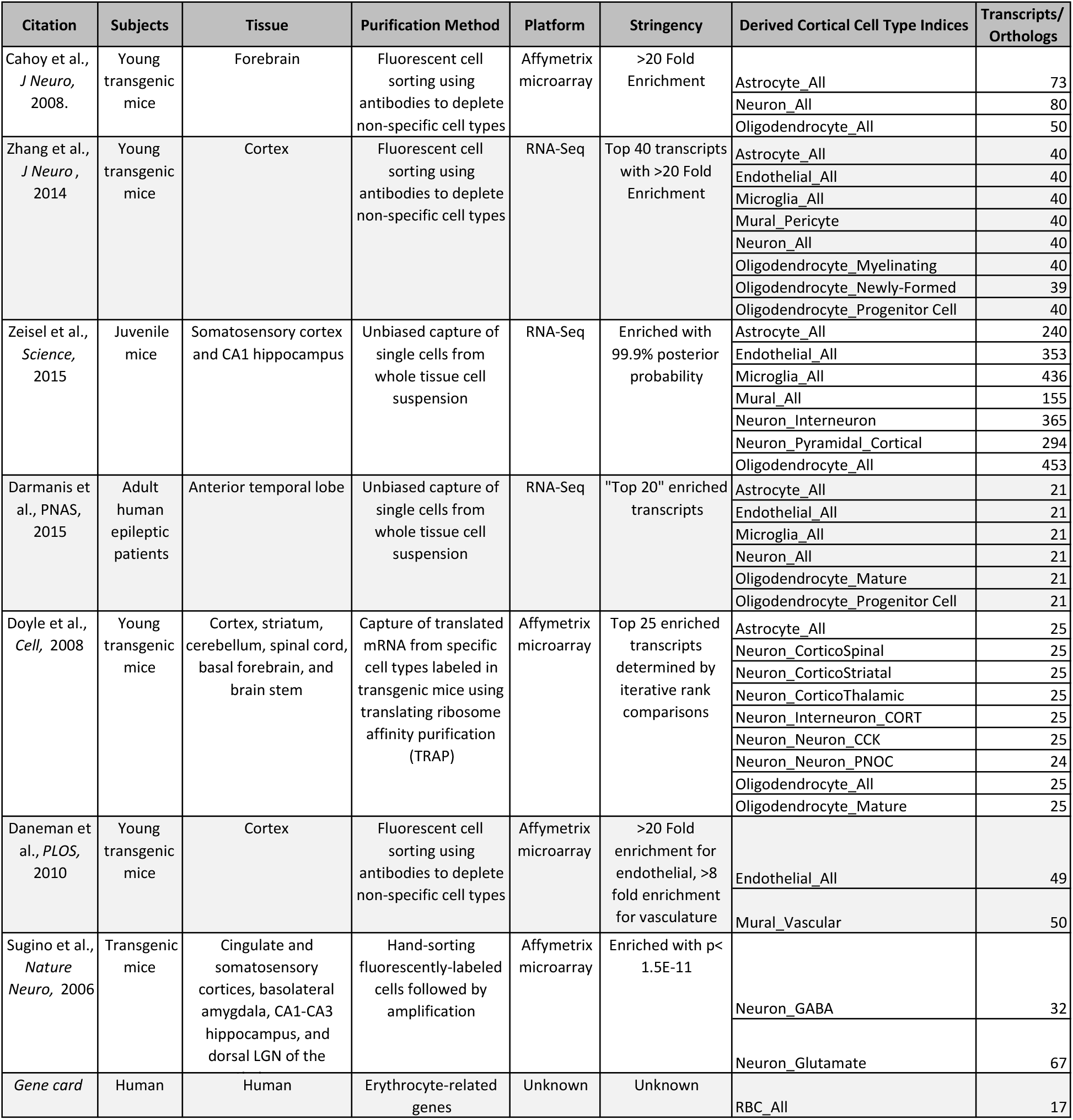
Thousands of transcripts have been identified as specifically-enriched in particular cortical cell types within published single-cell or purified cell type transcriptomic experiments. The experimental and statistical methods for determining whether a transcript was enriched in a cell type varied by publication, and included both RNA-Seq and microarray datasets.

### 2.2 “*BrainInABlender*”: Employing the Database of Cell Type Specific Transcripts to Predict Relative Cell Type Balance in Heterogenous Brain Samples

Next, we designed a method that uses the collective expression of cell type specific transcripts in brain tissue samples to predict the relative cell type balance of the samples (“BrainInABlender”). We specifically designed BrainInABlender to be compatible with large human brain transcriptional profiling datasets such as those used by our research consortium (Pritzker) and the Allen Brain Institute, which may lack full information about relative levels of expression within individual samples due to the extensive normalization procedures used to combine data across batches or platforms. We have made our method publicly-available in the form of a downloadable R package (https://github.com/hagenaue/BrainInABlender).

In brief, BrainInABlender extracts the data from transcriptional profiling datasets that represent genes identified in our database as having cell type specific expression in the brain (as curated by official gene symbol). Prior to application of our method, the dataset should be in the format of expression-level summary data (RNA-Seq: gene-level summary - CPM, RPKM or TPM; microarray: probe or probeset summary), and should have received at least some basic preprocessing, including log(2) transformation, normalization to eliminate technical variation, and standard quality control. Within BrainInABlender, these data are then centered and scaled across samples (mean=0, sd=1) to prevent transcripts with more variable signal from exerting disproportionate influence on the results. Then, if necessary, the normalized data from all transcripts representing the same gene are averaged for each sample and re-scaled. Finally, for each sample, these values are averaged across the genes identified as having expression specific to a particular cell type for each reference publication included in the database of cell type specific transcripts. This creates 38 cell type signatures derived from the cell type specific genes identified by the eight publications (“Cell Type Indices”), each of which predicts the relative content for one of the 10 primary cell types in our brain samples (**Figure 1**).

**Figure 1.**
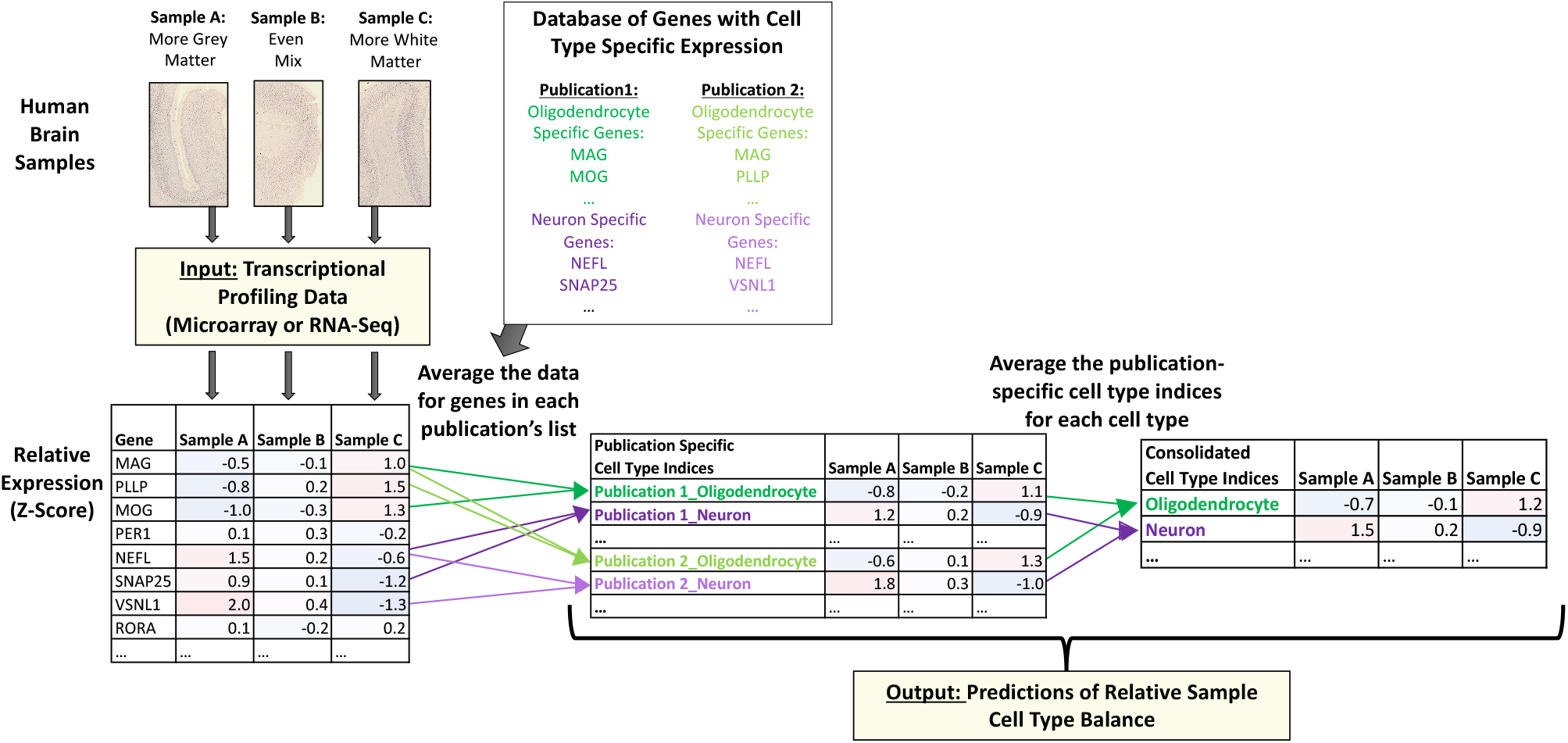
Predicting the relative cell type balance in human brain samples using genes previously-identified as having cell type specific expression. Within macro-dissected brain tissue samples, variable cell type balance is likely to influence the pattern of gene expression. To estimate this variability, we extracted the data for genes that had been previously identified as having cell type specific expression in previous publications (“Database of genes with cell type specific expression”, **Table 1**) and then averaged across the transcripts identified as specific to a particular cell type for each reference publication in our database to create 38 different "Cell Type Indices" that predicted relative cell content in each of the brain samples. Then, for many analyses in our paper, these publication-specific cell type indices were averaged within their cell type category to produce consolidated cell type indices representing each of the 10 primary cell types in the human cortex.

Later, during validation analyses (***Suppl. Section 7.2***), we found substantial support for simply averaging these 38 publication-specific cell type indices within each of the primary categories to produce ten consolidated primary cell-type indices for each sample. To perform this consolidation, we also removed any transcripts that were identified as “cell type specific” to multiple primary cell type categories (**Suppl. Figure 3**). These consolidated indices are included as output from BrainInABlender.

Please note that our method was specifically designed to tackle challenges present in our microarray data, but we later discovered that it bears some resemblance to the existing method of Population Specific Expression Analysis (PSEA, (10–12)). A more detailed discussion of the similarities and differences between the techniques can be found in ***Suppl. Section 7.2.2.***

### 2.3 Validation of Relative Cell Content Predictions

We initially validated our method using publicly-available datasets from purified cortical cell types (RNA-seq datasets GSE52564 and GSE6783), artificial mixtures of cells produced *in silico* by sampling from within these datasets, and microarray data from samples containing artificially-generated mixtures of cultured cells from P1 pups (Affymetrix Rat Genome 230 2.0 Array dataset GSE19380; further detail: ***Suppl. Sections 7.1.1-7.1.2***). For each of these analyses, we examined the correlation between the cell type indices outputted by BrainInABlender and the documented cell content of the samples.

Next, we wanted to see whether the cell content predictions produced by BrainInABlender could also correctly reflect relative cell type balance in human post-mortem samples. To test this, we applied our method to a large human post-mortem Agilent microarray dataset (841 samples) spanning 160 cortical and subcortical brain regions from the Allen Brain Atlas (http://human.brain-map.org/microarray/search, December 2015, (25)). This dataset was derived from high-quality tissue (absence of neuropathology, pH>6.7, post-mortem interval<31 hrs, RIN>5.5) from 6 human subjects (26). The tissue samples were collected using a mixture of block dissection and laser capture microscopy (27). After applying BrainInABlender, we compared the outputted cell type index results between selected brain regions known to contain relatively more (+) or less (-) of a particular cell type using Welch’s t-test (further detail: ***Suppl. Section 7.1.4**)*.

### 2.4 Predicting Relative Cell Content in Transcriptomic Data from Macro-Dissected Human Cortical Tissue from Psychiatric Subjects

Next, we profiled cell type specific gene expression in several large psychiatric human brain microarray datasets. The first was a large Pritzker Consortium Affymetrix U133A microarray dataset derived from high-quality human post-mortem dorsolateral prefrontal cortex samples (final n=157 subjects), including tissue from subjects without a psychiatric or neurological diagnosis (“Controls”, n=71), or diagnosed with Major Depressive Disorder (“MDD”, n=40), Bipolar Disorder (“BP”, n=24), or Schizophrenia (“Schiz”, n= 22). The severity and duration of physiological stress at the time of death was represented by an agonal factor score for each subject (ranging from 0-4, with 4 representing severe physiological stress (28,29)). We measured the pH of cerebellar tissue to indicate the extent of oxygen deprivation experienced around the time of death (28,29) and calculated the interval between the estimated time of death and the freezing of the brain tissue (the postmortem interval or PMI) using coroner records. Our current analyses began with subject-level summary gene expression data (GSE92538).

We determined the replicability of our results using three smaller publicly-available post-mortem human cortical Affymetrix U133Plus2 microarray datasets (GSE53987 (30), GSE21935 (31), GSE21138 (32), **Table 2.**). These datasets were selected because they included both psychiatric and control samples, and provided pH, PMI, age, and gender in the demographic information on the GEO website (https://www.ncbi.nlm.nih.gov/geo/). To control for technical variation, the sample processing batches were estimated using the microarray chip scan dates extracted from the .CEL files and RNA degradation was estimated using the R package AffyRNADegradation (33).

Finally, we also explored replicability within the recently-released large CommonMind Consortium (CMC) human dorsolateral prefrontal cortex RNA-seq dataset (34); downloaded from the CommonMind Consortium Knowledge Portal (https://www.synapse.org/CMC; final n=514 subjects). We predicted the relative cell type content of these samples using a newer version of BrainInABlender (v2) which excluded a few of the weaker cell type specific gene sets (15).

In general, the full preprocessing methods for these datasets can be found in ***Suppl. Section 7.1.*** The code for all analyses in the paper can be found at https://github.com/hagenaue/ and https://github.com/aschulmann/CMC_celltype_index.

**Table 2.**
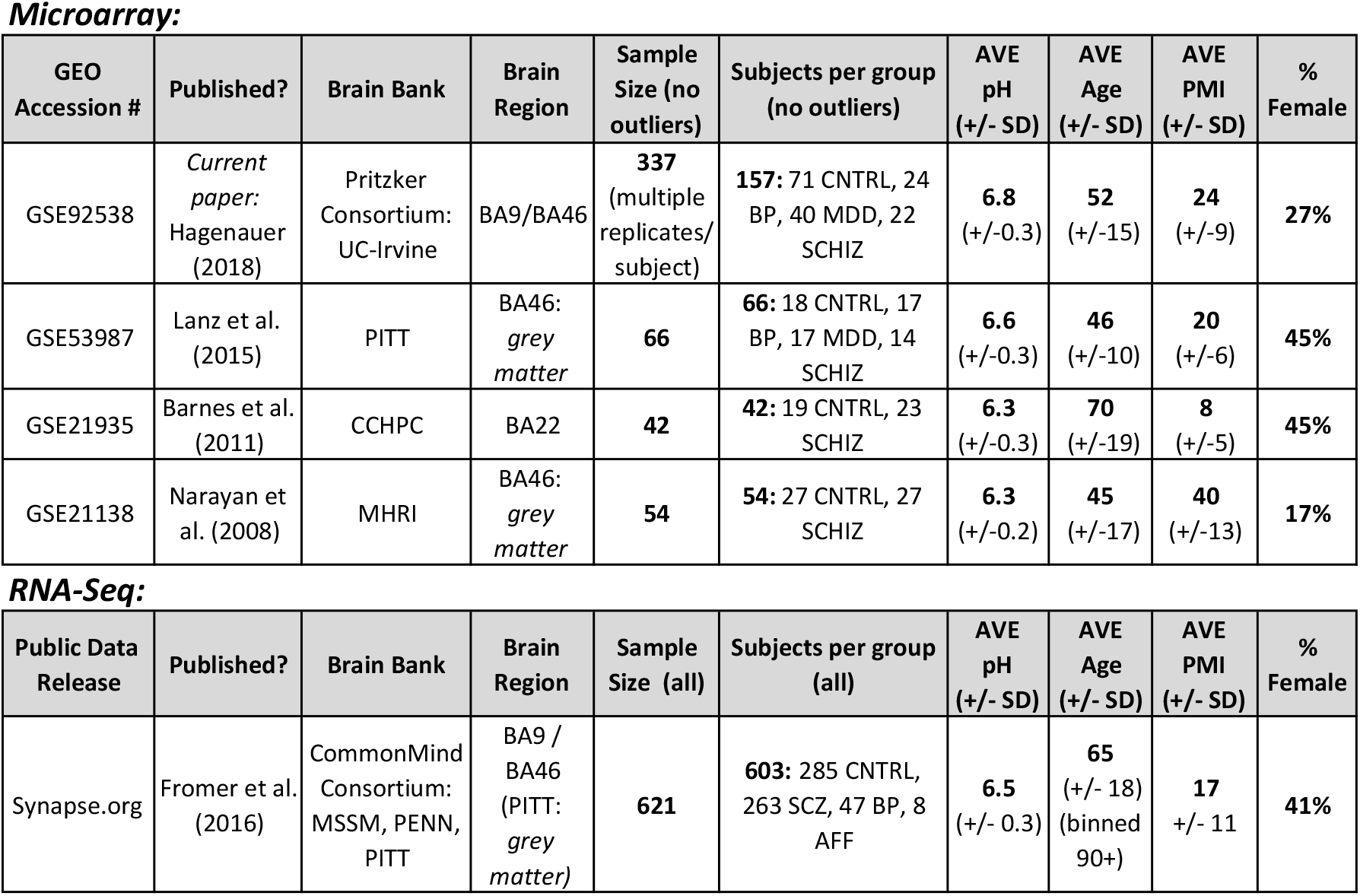
We examined the pattern of cell-type specific gene expression in five post-mortem human cortical tissue datasets that included samples from subjects with psychiatric illness. Abbreviations: CTRL: control, BP: Bipolar Disorder, MDD: Major Depressive Disorder, SCHIZ: Schizophrenia, GEO: Gene Expression Omnibus, BA: Brodmann’s Area, PMI: Post-mortem interval, SD: Standard Deviation, Brain Banks: UC-Irvine (University of California – Irvine), PITT (University of Pittsburgh), CCHPC (Charing Cross Hospital Prospective Collection), MSSM (Mount Sinai Icahn School of Medicine), MHRI (Mental Health Research Institute Australia), PENN (University of Pennsylvania)

**Microarray:**
| GEO Accession # | Published? | Brain Bank | Brain Region | Sample Size (no outliers) | Subjects per group (no outliers) | AVE pH (+/- SD) | AVE Age (+/- SD) | AVE PMI (+/- SD) | % Female |
| --- | --- | --- | --- | --- | --- | --- | --- | --- | --- |
| GSE92538 | *Current paper:*<br>Hagenauer (2018) | Pritzker Consortium:<br>UC-Irvine | BA9/BA46 | 337<br>(multiple replicates/<br>subject) | 157: 71 CNTRL, 24 BP, 40 MDD, 22 SCHIZ | 6.8<br>(+/-0.3) | 52<br>(+/-15) | 24<br>(+/-9) | 27% |
| GSE53987 | Lanz et al. (2015) | PITT | BA46:<br>*grey matter* | 66 | 66: 18 CNTRL, 17 BP, 17 MDD, 14 SCHIZ | 6.6<br>(+/-0.3) | 46<br>(+/-10) | 20<br>(+/-6) | 45% |
| GSE21935 | Barnes et al. (2011) | CCHPC | BA22 | 42 | 42: 19 CNTRL, 23 SCHIZ | 6.3<br>(+/-0.3) | 70<br>(+/-19) | 8<br>(+/-5) | 45% |
| GSE21138 | Narayan et al. (2008) | MHRI | BA46:<br>*grey matter* | 54 | 54: 27 CNTRL, 27 SCHIZ | 6.3<br>(+/-0.2) | 45<br>(+/-17) | 40<br>(+/-13) | 17% |

**Table 2.** We examined the pattern of cell-type specific gene expression in five post-mortem human cortical tissue datasets that included samples from subjects with psychiatric illness.
| Public Data Release | Published? | Brain Bank | Brain Region | Sample Size (all) | Subjects per group (all) | AVE pH (+/- SD) | AVE Age (+/- SD) | AVE PMI (+/- SD) | % Female |
| --- | --- | --- | --- | --- | --- | --- | --- | --- | --- |
| Synapse.org | Fromer et al. (2016) | CommonMind Consortium:<br>MSSM, PENN, PITT | BA9 / BA46<br>(PITT: *grey matter*) | 621 | 603: 285 CNTRL, 263 SCZ, 47 BP, 8 AFF | 6.5<br>(+/- 0.3) | 65<br>(+/- 18)<br>(binned 90+) | 17<br>+/- 11 | 41% |

### 3.5 Examining the Relationship Between Predicted Cell Content Derived from Transcriptional Profiling Data and Clinical/Biological Variables

We next set out to observe the relationship between the predicted cell content of our samples and a variety of medically-relevant subject variables. To perform this analysis, we first examined the relationship between seven relevant subject variables and each of the ten consolidated cell type indices in the Pritzker prefrontal cortex dataset using a linear regression model that allowed us to simultaneously control for other likely confounding variables:

***Equation 1:***

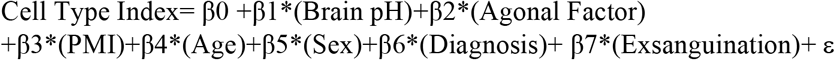

We then examined the replicability of these relationships using data from the three smaller publicly-available human post-mortem microarray datasets (GSE53987, GSE21935, GSE21138). For these datasets, we initially lacked detailed information about manner of death (agonal factor and exsanguination), but were able to control for technical variation within the model using statistical estimates of RNA degradation and batch (scan date):

***Equation 2:***

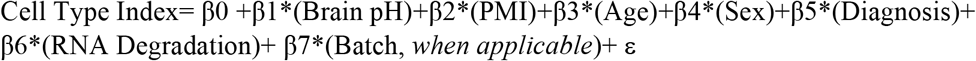

We evaluated replicability by performing a meta-analysis for each variable and cell type combination across the four microarray datasets. To do this, we applied random effects modeling to the respective betas and accompanying sampling variance derived from each dataset using the *rma.mv()* function within the *metafor* package (35). P-values were corrected for multiple comparisons following the Benjamini-Hochberg method (FDR or q-value; (36)).

Finally, we characterized these relationships in the large CMC RNA-seq dataset. For this dataset, we had some information about manner of death but lacked knowledge of agonal factor or exsanguination. We controlled for technical variation due to dissection site (institution) and RNA degradation (RIN):

***Equation 3:***

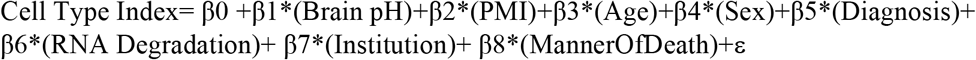

### 3.6 Characterizing Psychiatric Gene Expression using Differential Expression Models that Include Either Standard Co-variates or Cell Type Indices

To determine whether controlling for variability in cell type balance in the dataset could improve our ability to detect differential expression related to psychiatric illness, we compared differential expression results within the human psychiatric datasets that were derived from linear regression models of increasing complexity, including a simple base model containing just the variable of interest (“Model 1”), a standard model controlling for traditional co-variates (“Model 2”), and a model controlling for traditional co-variates as well as each of the cell type indices (“Model 5”). We also used two reduced models that only included the most prevalent cell types (Astrocyte, Microglia, Oligodendrocyte, Neuron_Interneuron, Neuron_Projection; (21)) to avoid issues with multicollinearity. The first of these models included traditional co-variates (“Model 4”), whereas the second model excluded them (“Model 3”) (**Equation 4**).

**Equation 4: A model of gene expression for each dataset, colored to illustrate the subcomponents evaluated during our model comparison (#M1-M5).** The base model (intercept and variable of interest) is presented in green, traditional subject variable covariates are blue, the cell type indices for the most prevalent cell types are red, and the remaining cell type indices are purple. Model components unique to each dataset are underlined.

***The Pritzker microarray dataset:***

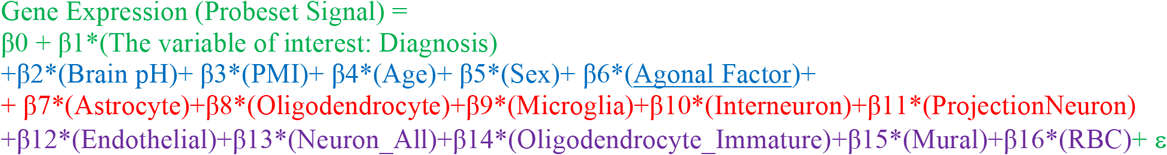

***The CMC RNA-Seq dataset:***

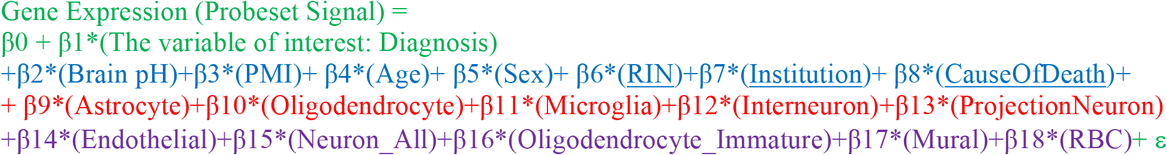

***The smaller microarray datasets (GSE53987, GSE21935, GSE21138):***

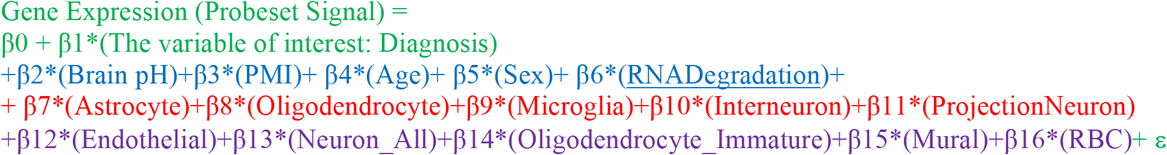

### 3.7 Functional Ontology with Cell Type Specific Gene Sets

We ran a series of analyses to evaluate how well we could distinguish between changes in cell type balance in the tissue and changes in cell type specific functions. First, as a case study, we specifically examined the relationship between age and the functional annotation for genes found in the Neuron_All index in more depth. To do this, we evaluated the relationship between age and gene expression in the Pritzker dataset using a standard model that controlled for traditional confounds (“Model 2”) using the signal data for all probesets in the dataset. We used “DAVID: Functional Annotation Tool” (//david.ncifcrf.gov/summary.jsp, (37,38) to identify the functional clusters that were overrepresented by the genes included in our neuronal cell type index (using the full HT-U133A chip as background), and then determined the average effect of age (beta) for the genes included in each of the 240 functional clusters. These functional clusters overrepresented dendritic/axonal related functions, so for a follow-up analysis, in a manner that was blind to the results, we subsetted the results into 29 functional clusters that were clearly related to dendritic/axonal functions and 41 functional clusters that seemed distinctly unrelated to dendritic/axonal functions (**Suppl. Table 4**) and compared the average effect of age in these two subsets using a Welch’s t-test.

In the next analysis, we decided to make the process of differentiating between altered cell type-specific functions and relative cell type balance more efficient. We used our cell type specific gene lists to construct gene sets in a file format (.gmt) compatible with the popular tool Gene Set Enrichment Analysis (GSEA, (39,40)) and combined them with two other commonly-used gene set collections from the molecular signatures database (MSigDB: http://software.broadinstitute.org/gsea/msigdb/index.jsp, downloaded 09/2017, “C2: Curated Gene Sets” and “C5: GO Gene Sets”, **Suppl. Table 5**). Then we tested the utility of incorporating our new gene sets into GSEA (fGSEA: (41)) using the ranked results (betas) for the relationship between each subject variable and each probeset in the Pritzker dataset (as evaluated using a standard model: “Model 2”).

## 3. Results & Discussion

### 3.1 Validation of Relative Cell Content Predictions

#### Validation Using Datasets Derived from Purified or Cultured Cells

We initially validated our method using publicly-available datasets from purified cell types (datasets GSE52564 and GSE6783; (2,18) and *in silico* derived mixtures and found that the statistical cell type indices easily predicted the cell type identities of the samples (***Suppl. Section 7.2.10***). Therefore, as further validation, we determined whether relative cell type balance could be accurately deciphered from microarray data for samples containing artificially-generated mixtures of cultured cells (GSE19380; (12)). We found that the consolidated cell type indices produced by BrainInABlender strongly correlated with the actual percentage of cells of a particular type included in the artificial mixtures (**Figure 2**, Neuron% vs. Neuron_All Index: R^2^=0.93, p=1.54e-15, Astrocyte% vs. Astrocyte Index: R^2^=0.77, p=5.05e-09, Microglia% vs. Microglia Index: R^2^=0.64, p=8.2e-07), although we found that the cell type index for immature oligodendrocytes better predicted the percentage of cultured oligodendrocytes in the samples than the cell type index for mature oligodendrocytes (Mature: R^2^=0.45, p=0.000179, Immature: R^2^=0.81, p=4.14e-10). We believe this discrepancy is likely to reflect the specific cell culture conditions used in the original admixture experiment. Notably, the relationship between the consolidated cell type indices and the actual percentage of each cell type included in the artificial mixtures was approximately linear, despite the use of log(2)-transformed expression data.

**Figure 2.**
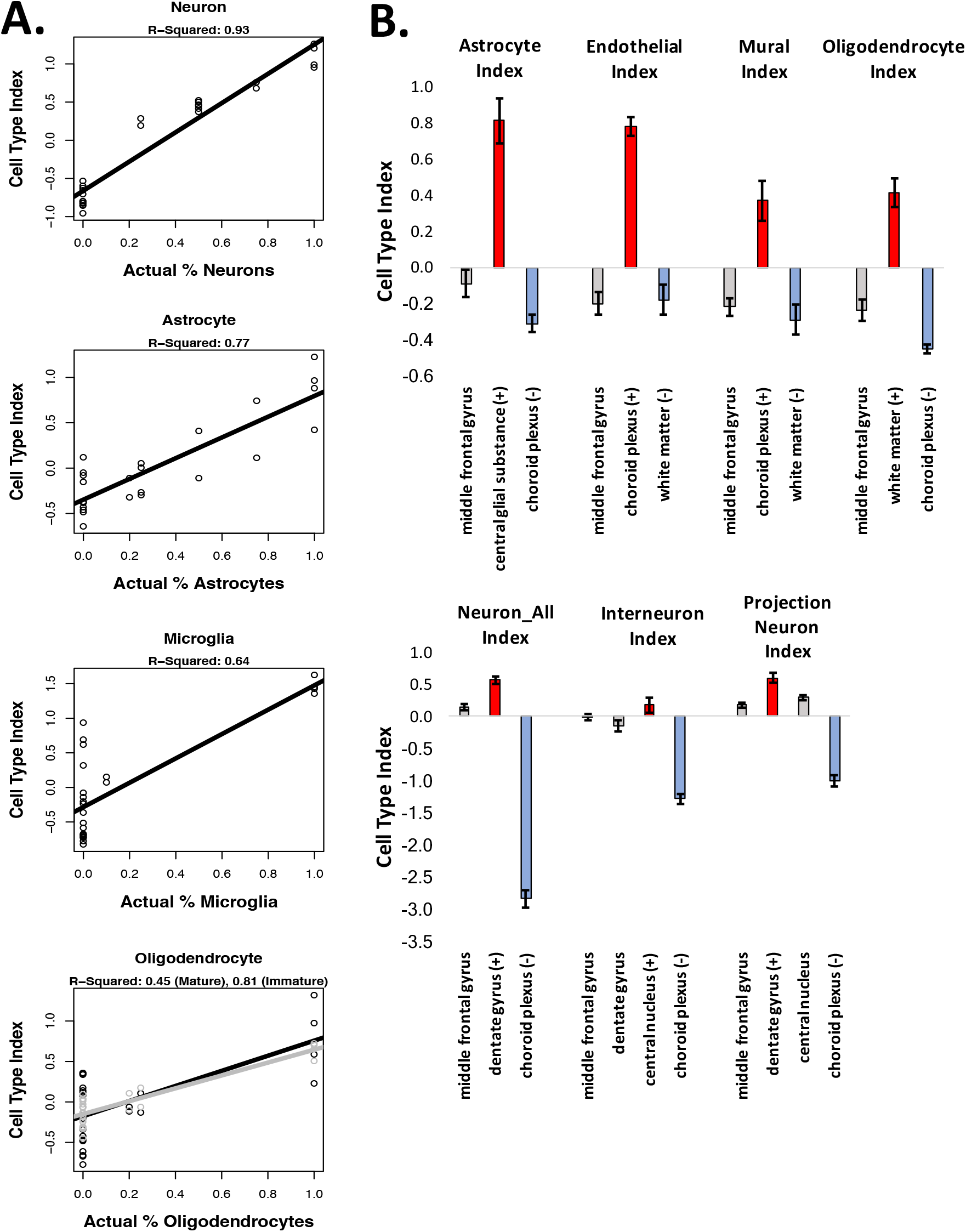
Validation of Relative Cell Content Predictions. A) Using a microarray dataset derived from samples that contained artificially-generated mixtures of cultured cells (GSE19380; (12)), we found that our relative cell content predictions (“cell type indices”) closely reflected actual known content. However, note that the numeric values for the cell type indices do not convey an absolute proportion of cells of a particular type in the sample - simply whether a sample contains relatively more or less of the cell type of interest in comparison to other samples in the dataset. B) Our cell type indices also easily differentiated human post-mortem samples derived from brain regions that are known to contain relatively more (+, red) or less (-, blue) of the targeted cell type of interest (all p<0.007). Results from the middle frontal gyrus are included for comparison, since the rest of the paper primarily focuses on prefrontal cortical data. (Bars: average +/-SE).

#### Validation Using a Dataset Derived from Human Post-Mortem Tissue

Next, we wanted to see whether the cell content predictions produced by BrainInABlender correctly reflected relative cell type balance in human post-mortem samples. To test this, we applied our method to a large cross-regional human post-mortem microarray dataset (25), and extracted the results for a selection of brain regions that are known to contain relatively more (+) or less (-) of particular cell types (the results for other regions can be found in **Suppl. Table 2**). The results clearly indicated that our cell type analyses could identify well-established differences in cell type balance across brain regions (**Figure 2**, (+) region vs. (-) region for all cell types: *p*<0.007, Cohen’s *d*>3.2). The choroid plexus had elevated gene expression specific to vasculature (endothelial cells, mural cells, (42)). The corpus callosum and cingulum bundle showed an enrichment of oligodendrocyte-specific gene expression (42). The central glial substance was enriched with gene expression specific to glia and support cells, especially astrocytes. The dentate gyrus, which contains densely-packed glutamatergic granule cells (43), was enriched for gene expression specific to projection neurons. The highly GABA-ergic central nucleus of the amygdala (44) had a slight enrichment of gene expression specific to interneurons. These results provide fundamental validation that our methodology can accurately predict relative cell type balance in human post-mortem samples. Moreover, these results suggest that the cell type indices are capable of generally tracking their respective cell types in subcortical structures, despite the dependency of our method on cell type specific gene lists derived from the forebrain and cortex.

### 3.8 Inferred Cell Type Composition Explains a Large Percentage of the Sample-Sample Variability in Transcriptomic Data from Macro-Dissected Human Cortical Tissue

Using principal components analysis we found that the primary gradients of gene expression variation across samples in all four of the cortical transcriptomic datasets strongly correlated with our estimates of cell type balance. For example, while analyzing the Pritzker microarray dataset, we found that the first principal component (PC1), which encompassed 23% of the variation in gene expression across samples in the dataset, spanned from samples with high predicted support cell content to samples with high predicted neuronal content. Therefore, a large percentage of the variation in PC1 (91%) was accounted for by an average of the astrocyte and endothelial indices (p=2.2e-82, with a respective R^2^ of 0.80 and 0.75 for each index analyzed separately) or by the general neuron index (p=6.3e-32, R^2^=0.59). The second notable gradient in the dataset (PC2) encompassed 12% of the variation overall, and spanned samples with high predicted projection neuron content to samples with high predicted oligodendrocyte content (with a respective R^2^ of 0.62 and 0.42, and p-values of p=8.5e-35 and p=8.7e-20).

To confirm that the strong relationship between the top principal components of variation and our cell type indices did not originate artificially due to cell type specific genes representing a large percentage of the most highly variable transcripts in the dataset, we repeated the principal components analysis after excluding all cell type specific transcripts from the dataset and still found these strong correlations (**Figure 3;** PC1 vs. average astrocyte/endothelial index: R^2^=0.89, p=1.1e-76; PC2 vs. projection neuron index: R^2^=0.65, p=1.6e-37). Indeed, individual cell type indices still better accounted for the main principal components of variation in the microarray data than *all other major subject variables combined* (pH, Agonal Factor, PMI, Age, Gender, Diagnosis; PC1: R^2^=0.416, PC2: R^2^=0.203). Similarly, when examining the data for individual probesets, a linear model that included just the six subject variables (**Equation 4**) accounted for an average of only 12% of the variation (R^2^, Adj.R^2^=0.0692), whereas a linear model including the astrocyte and projection neuron indices alone accounted for 17% (R^2^, Adj.R^2^=0.156) and a linear model including all 10 cell types accounted for an average of 30% (R^2^, Adj.R^2^=0.255), almost one third of the variation present in the data for any particular probeset. Therefore, a large percentage of the genes in our dataset seemed to be preferentially expressed in relationship to particular cell types, even if their expression was not defined as strictly cell type specific in our database.

**Figure 3.**
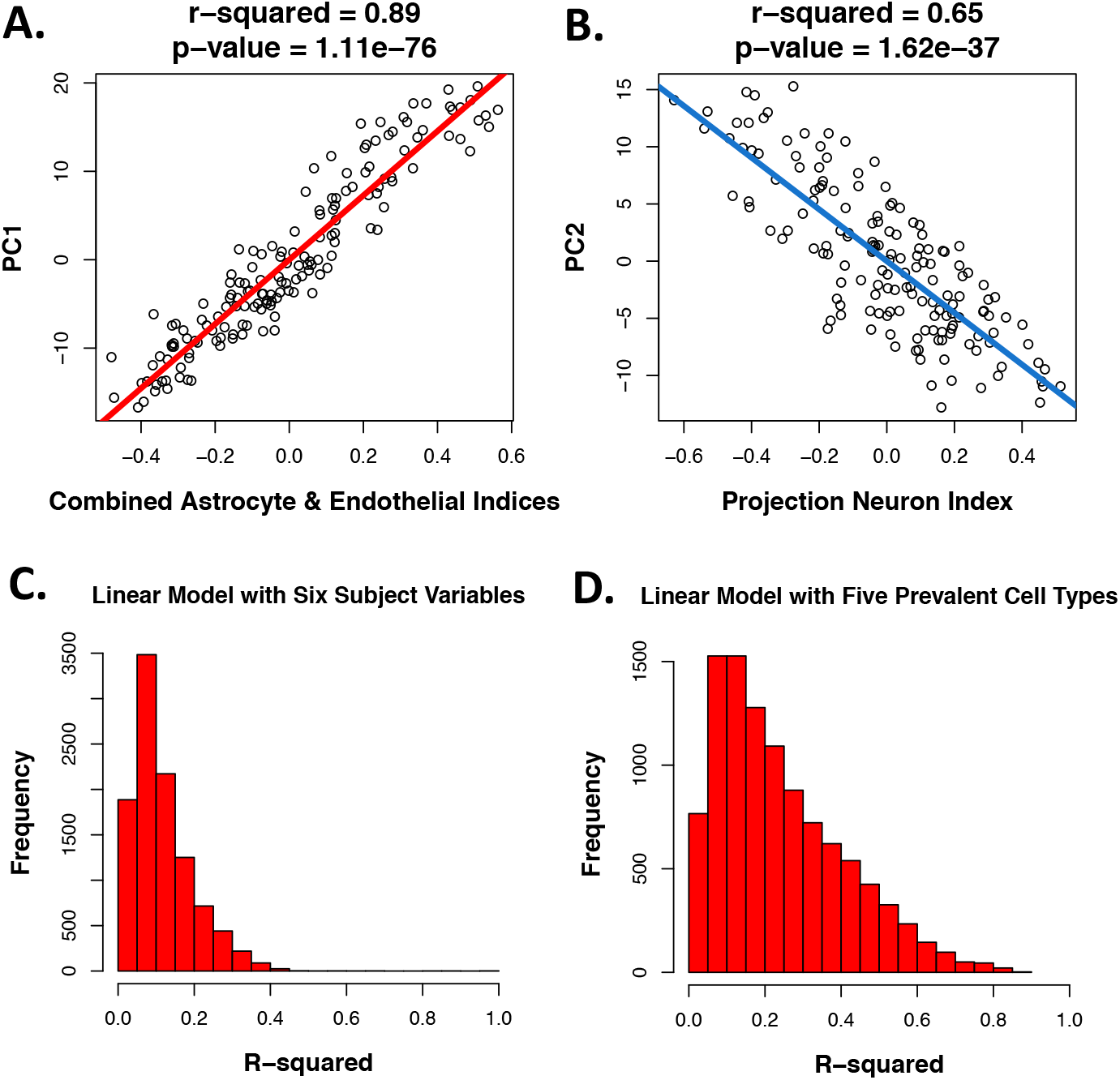
Cell content predictions explain a large percentage of the variability in microarray data derived from the human cortex. As an example, within the Pritzker dataset, even after excluding all data from genes identified as cell type specific in our database, **A)** the first principal component of variation (PC1) was strongly correlated with predicted “support cell” content in the samples (the average of the astrocyte and endothelial indices). **B)** PC2 was strongly correlated with predicted projection neuron content. Likewise, when applying a linear model to the data for each probeset, the R^2^ values for each probeset (illustrated in the histogram) tended to be much smaller when using a model that included **C)** only the six subject variables, versus **D**) only the five most prevelant cortical cell types.

Within the other four human cortical tissue datasets, the relationships between the top principal components of variation and the consolidated cell type indices were similarly strong (***Suppl. Section 3.8***), despite the fact that these datasets had received less preprocessing to remove the effects of technical variation. These results indicated that accounting for cell type balance is important for the interpretation of post-mortem human brain transcriptomic data and might improve the signal-to-noise ratio in analyses aimed at identifying psychiatric risk genes.

### 3.9 Cell Content Predictions Derived from Transcriptional Profiling Data Match Known Relationships Between Clinical/Biological Variables and Brain Tissue Cell Content

We next set out to observe the relationship between the predicted cell content of our samples and a variety of medically-relevant subject variables. This analysis uncovered many relationships that had been previously-identified using other paradigms or animal models (**Figure 4, Suppl. Table 3**).

**Figure 4.**
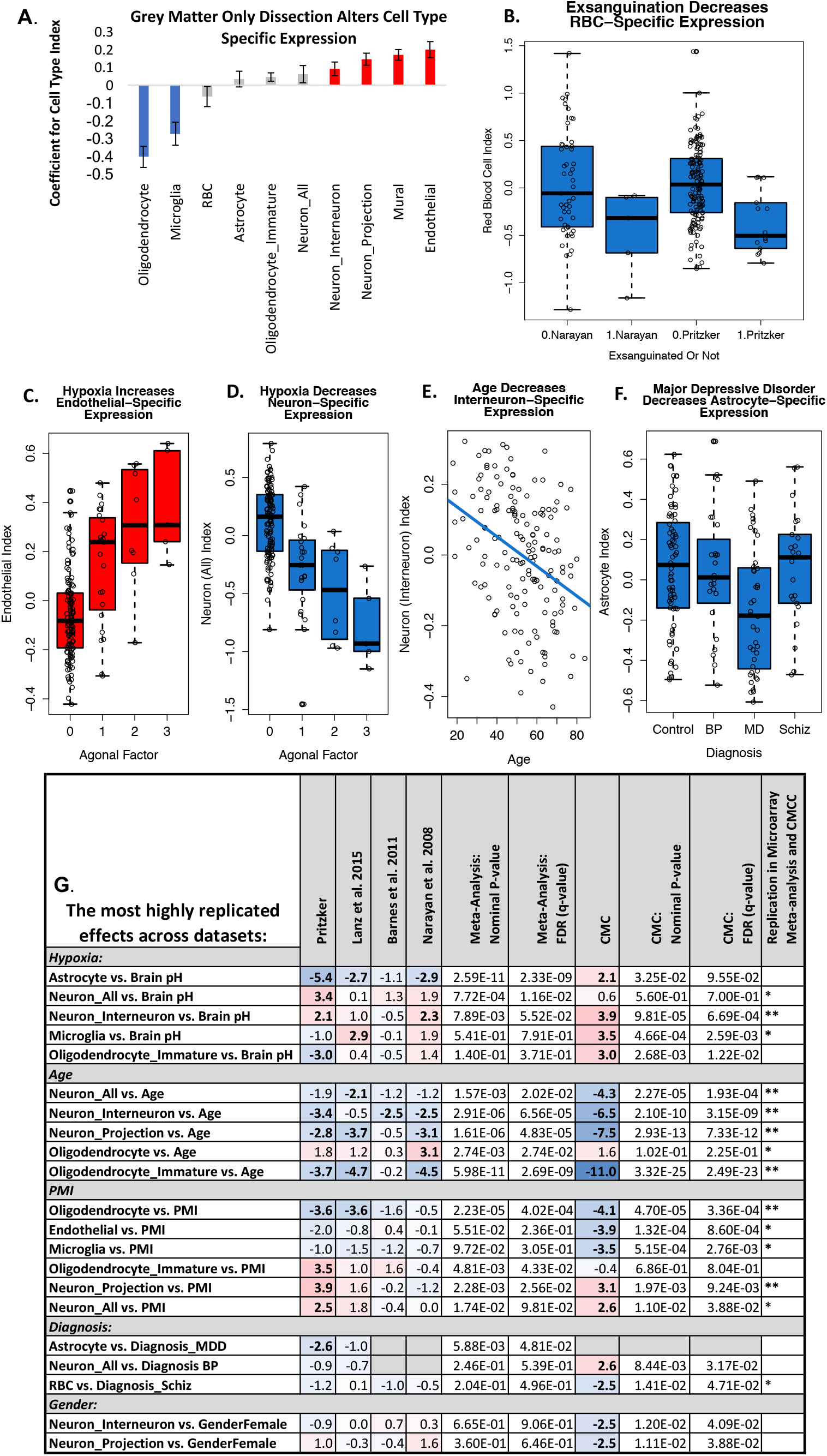
Cell content predictions derived from microarray data match known relationships between subject variables and brain tissue cell content. Boxplots represent the median and interquartile range, with whiskers illustrating either the full range of the data or 1.5x the interquartile range. **A.** Within the CMC dataset, cortical tissue samples that were dissected to only contain gray matter (PITT) show lower predicted oligodendrocyte and microglia content and more neurons and vasculature (bars: ß+/-SE, red/blue: p<0.05). **B.** Subjects who died in a manner that involved exsanguination had a notably low red blood cell index in both the Pritzker (p=0.00056) and Narayan et al. datasets (p=0.052*trend). **C.** The presence of prolonged hypoxia around the time of death, as indicated by high agonal factor score, was associated with a large increase in the endothelial cell index (p=2.85e-07) matching previous demonstrations of cerebral angiogenesis, activation, and proliferation in low oxygen environments (45). **D.** High agonal factor was also associated with a clear decrease in neuronal indices (p=3.58e-09) mirroring the vulnerability of neurons to low oxygen (46). **E.** Age was associated with a decrease in the neuronal indices (p= 0.000956) which fits known decreases in gray matter density in the frontal cortex in aging humans (47). **F**. Major Depressive Disorder was associated with a moderate decrease in astrocyte index (p= 0.0118), which fits what has been observed morphometrically (48). **G.** The most highly-replicated relationships between subject variables and predicted cortical cell content across all five of the post-mortem human datasets. Provided in the table are the T-stats for the effects (red=upregulation, blue=downregulation), derived from a larger linear model controlling for confounds **(Equation 1, Equation 2, Equation 3**), as well as the nominal p-values from the meta-analysis of the results across the four microarray studies, and p-values following multiple-comparisons correction (q-value). Only effects that had a q<0.05 in either our meta-analysis or the large CMC RNA-Seq dataset are included in the table. Asterisks denote effects that had consistent directionality in the meta-analysis and CMC dataset (*) or consistent directionality and q<0.05 in both datasets (**). Please note that lower pH and higher agonal factor are both indicators of greater hypoxia prior to death, but have an inverted relationship and therefore show opposing relationships with the cell type indices.

#### Dissection

First, as a proof of principle, we were able to clearly observe dissection differences between institutions within the large CMC RNA-Seq dataset, with samples from University of Pittsburgh having a predicted relative cell type balance that closely matched what would be expected due to their gray matter only dissection method (Oligodendrocyte: ß =-0.404, p=2.42e-11; Microglia: β=-0.274, p=3.06e-05; Neuron_Interneuron: β=0.0916, p=0.0161; Neuron_Projection: β=0.145, p=2.31e-05; Mural: β=0.170, p=2.14e-08; Endothelial: β=0.200, p=1.12e-05). In contrast, samples from University of Pennsylvania were associated with lower predicted cell content related to vasculature (Endothelial: β=-0.255, p=4.01-04; Mural: β=-0.168, p=4.59e-04; Astrocyte: β=-0.189, p=7.47e-03).

#### Manner of Death

Predicted cell type content was also closely related to manner of death. Within the Pritzker dataset we found that subjects who died in a manner that involved exsanguination had a notably low red blood cell index (β=-0.398; p=0.00056). Later, we were able replicate this result within GSE21138 using data from 5 subjects who were also likely to have died in a manner involving exsanguination (β=-0.516, p=0.052*\*trend*, manner of death reported in suppl. in *(32)*).

The presence of prolonged hypoxia around the time of death, as indicated by either low brain pH or high agonal factor score within the Pritzker dataset, was associated with a large increase in the endothelial cell index (Agonal Factor: β=0.118 p=2.85e-07; Brain pH: β=-0.210, p=0.0003) and astrocyte index (Brain pH: β=-0.437, p=2.26e-07; Agonal Factor: β=0.071, p=0.024), matching previous demonstrations of cerebral angiogenesis, endothelial and astrocyte activation and proliferation in low oxygen environments (45). Smaller increases were also seen in the mural index (Mural vs. Agonal Factor: β= 0.0493, p=0.0286). In contrast, prolonged hypoxia was associated with a clear decrease in all of the neuronal indices (Neuron_All vs. Agonal Factor: β=-0.242, p=3.58e-09; Neuron_All vs. Brain pH: β=0.334, p=0.000982; Neuron_Interneuron vs. Agonal Factor: β=-0.078, p=4.13e-05; Neuron_Interneuron vs. Brain pH: β=0.102, p=0.034; Neuron_Projection vs. Agonal Factor: β=-0.096, p= 0.000188), mirroring the notorious vulnerability of neurons to low oxygen (e.g., (46)).

These overall effects of hypoxia on predicted cell type balance replicated in the smaller human microarray post-mortem datasets (Astrocyte vs. Brain pH (meta-analysis: b=-0.459, p=2.59e-11): GSE21138: β=-0.856, p=0.00661, GSE53987: β=-0.461, p=0.00812, Neuron_All vs. Brain pH (meta-analysis: b= 0.245, p=7.72e-04), Neuron_Interneuron vs. Brain pH (meta-analysis: b=0.109, p=7.89e-03): GSE21138: β=0.381134, p=0.0277), despite lack of information about agonal factor, and partially replicated in the CMC human RNA-Seq dataset (Neuron_Interneuron vs. Brain pH: β=0.186, p=9.81e-05). In several datasets, we also found that prolonged hypoxia correlated with a decreased microglial index (Microglia vs. Brain pH: GSE53987: β=0.462, p=0.00603; CMC: β=0.286, p=4.66e-04).

#### Age

In the Pritzker dataset, age was associated with a moderate decrease in two of the neuronal indices (Neuron_Interneuron vs. Age: β=-0.00291, p=0.000956; Neuron_Projection Neuron vs. Age: β=-0.00336, p=0.00505) and this was strongly replicated in the large CMC RNA-Seq dataset (Neuron_All vs. Age: β=-0.00497, p=2.27e-05; Neuron_Projection Neuron vs. Age: β=-0.00612, p=2.93e-13; Neuron_Interneuron vs. Age: β=-0.00591, p=2.10e-10). A similar decrease in predicted neuronal content was seen in all three of the smaller human post-mortem datasets (Neuron_All vs. Age (meta-analysis: b=-0.00415, p=1.57e-03): GSE53987: β=-0.00722, p=0.0432, Neuron_Interneuron vs. Age (meta-analysis: b=-0.00335, p=2.91e-06): GSE21138: β=-0.00494, p=0.0173, GSE21935: β=-0.00506, p=0.0172, Neuron_Projection vs. Age (meta-analysis: b=-0.00449, p=1.61e-06): GSE53987: β=-0.0103, p=0.000497, GSE21138: β=-0.00763, p=0.00386). This result mirrors known decreases in gray matter density in the frontal cortex in aging humans (47), as well as age-related sub-region specific decreases in frontal neuron numbers in primates (49) and rats (50).

There was a consistent decrease in the immature oligodendrocyte index in relationship to age across datasets (Oligodendrocyte_Immature vs. Age (meta-analysis: b=-0.00514, p=5.98e-11): Pritzker: β=-0.00432, p=0.000354, GSE21138: β=-0.00721, p=5.73e-05, GSE53987: β=-0.00913, p=1.85e-05; CMC: β=-0.00621, p=3.32e-25), which seems intuitive, but actually contradicts animal studies on the topic (51). Since the validation of the immature oligodendrocyte index was relatively weak (***Suppl. Section 7.2***), this result should perhaps be considered with caution.

In some datasets, there also appeared to be an increase in the oligodendrocyte index with age (Oligodendrocyte vs. Age (meta-analysis: b=0.00343, p=2.74e-03): GSE21138, β=0.00957, p=0.00349) which, at initial face value, seems to contrast with well-replicated observations that frontal white matter decreases with age in human imaging studies (47,52,53). However, it is worth noting that several histological studies in aging primates suggest that brain regions that are experiencing demyelination with age actually show an *increasing* number of oligodendrocytes due to repair (51,54).

#### PMI

A prominent unexpected effect was a large decrease in the oligodendrocyte index with longer post-mortem interval (Oligodendrocyte vs. PMI (meta-analysis: b=-0.00764, p=2.23e-05): Pritzker: β=-0.00749, p=0.000474, GSE53987: β=-0.0318, p=0.000749; CMC: β=-0.00759, p=4.70e-05). Upon further investigation, we found a publication documenting a 52% decrease in the fractional anisotropy of white matter with 24 hrs post-mortem interval as detected by neuroimaging (55), but to our knowledge the topic is otherwise not well studied. These changes were paralleled by a decrease in the endothelial index (CMC: β=-0.00542, p=1.32e-04) and microglial index (CMC: β=-0.00710, p=5.15e-04) and increase in the immature oligodendrocyte index (Oligodendrocyte_Immature vs. PMI (meta-analysis: b=0.00353, p=4.81e-03): Pritzker: β=0.00635, p=0.000683) and neuronal indices (Neuron_All vs. PMI: Pritzker: β=0.006997, p=0.000982; CMC: β=0.00386, p=0.0110; Neuron_Projection vs. PMI (meta-analysis: b=0.00456, p=2.28e-03): Pritzker: β= 0.00708, p=1.64e-04; CMC: β=0.00331, p=0.00197). These results could arise from the zero-sum nature of transcriptomics analysis: due to the use of a standardized dissection size, RNA concentration, and data normalization, if there are large decreases in gene expression for one common variety of cell type (oligodendrocytes), then gene expression related to other cell types may appear to increase.

#### Psychiatric Diagnosis

Of most interest to us were potential changes in cell type balance in relation to psychiatric illness. In previous post-mortem morphometric studies, there was evidence of glial loss in the prefrontal cortex of subjects with Major Depressive Disorder, Bipolar Disorder, and Schizophrenia (reviewed in (56)). This decrease in glia, and particularly astrocytes, was replicated experimentally in animals exposed to chronic stress (57), and when induced pharmacologically, drove animals into a depressive-like condition (57). Replicating the results of (48), we observed a moderate decrease in astrocyte index in the prefrontal cortex of subjects with Major Depressive Disorder (meta-analysis: b=0.132, p=5.88e-03, Pritzker: β =-0.133, p=0.0118, **Figure 4 F**), but did not see similar changes in the brains of subjects with Bipolar Disorder or Schizophrenia.We also observed a decrease in red blood cell index in association with Schizophrenia (CMC: β=-0.104, p=0.0141) which is tempting to ascribe to reduced blood flow due to hypofrontality (58). This decrease in red blood cell content could also arise due to psychiatric subjects having an increased probability of dying a violent death, but the effect remained present when we controlled for exsanguination, and therefore is likely to be genuinely tied to the illness itself.

#### General Discussion

Overall, these results indicate that statistical predictions of the cell content of samples effectively capture many known biological changes in cell type balance, and imply that within both chronic (age, diagnosis) and acute conditions (agonal, PMI, pH) there is substantial influences upon the relative representation of different cell types.

The effect of hypoxia within our results is particularly worth discussing in greater depth. It has been acknowledged for a long time that exposure to a hypoxic environment prior to death has a huge impact on gene expression in human post-mortem brains (e.g., (28,29,59–61)). This impact on gene expression is so large that up until recently the primary principal component of variation (PC1) in our data was assumed to represent the degree of hypoxia, and was sometimes even removed before performing diagnosis-related analyses (e.g., (62)). These large effects of hypoxia on gene expression were hypothesized to be partially mediated by neuronal necrosis (63) and lactic acidosis (60). However, the magnitude of the effect of hypoxia was still puzzling, especially when compared to the much more moderate effects of post-mortem interval (even when ranging from 8-40+ hrs). Our current analysis provides an explanation for this discrepancy, since it is clear from our results that the brains of our subjects are *actively compensating* for a hypoxic environment prior to death by altering the balance or overall transcriptional activity of support cells and neurons. The differential effects of hypoxia on neurons and glial cells have been studied since the 1960’s (64), but to our knowledge this is the first time that anyone has related the large effects of hypoxia in post-mortem transcriptomic data to a corresponding upregulation in the transcriptional activity of vascular cell types (45).

This connection is important for understanding why results associating gene expression and psychiatric illness in human post-mortem tissue sometimes do not replicate. If a study contains mostly tissue from individuals who experienced greater hypoxia before death (e.g., hospital care with artificial respiration or coma), then differential expression analyses are likely to inadvertently focus on neuropsychiatric effects in support cell types, whereas a study that mostly contains tissue from individuals who died a fast death (e.g., myocardial infarction) will emphasize the neuropsychiatric effects in neurons. That said, although both indicators of perimortem hypoxia (agonal factor and pH) showed similar strong relationships with cell type balance, we recommend caution when interpreting the relationship between pH and cell type in tissue from psychiatric subjects, as pH can indicate other biological changes besides hypoxia. For example, there are small consistent decreases in pH associated with Bipolar Disorder even in live subjects (65–67) and metabolic changes associated with pH are theorized to play an important role in Schizophrenia (61). Therefore, the relationship between pH and cell type balance may be partially driven by a third variable (psychiatric illness or treatment). It is also possible that changes in tissue cell content could cause a change in pH (68).

### 3.10 It is Difficult to Discriminate Between Changes in Cell Type Balance and Cell-Type Specific Function

Gray matter density has been shown to decrease in the frontal cortex in aging humans (47), and frontal neuron numbers decrease in specific subregions in aging primates (49) and rats (50). However, many scientists would argue that age-related decreases in gray matter are primarily driven by synaptic atrophy instead of decreased cell number (69). This raised the question of whether the decline that we saw in neuronal cell indices with age was being largely driven by the enrichment of genes related to synaptic function in the index. More generally, it raised the question of how well cell type indices could discriminate changes in cell number from changes in cell-type function.

We examined this question using two methods. First, as a case study, we specifically examined the relationship between age and the functional annotation for genes found in the Neuron_All index in more depth. We found that transcripts from functional clusters that seemed distinctly unrelated to dendritic/axonal functions still showed an average decrease in expression with age (T(40)=-2.7566, p=0.008756), but this decrease was larger for transcripts clearly associated with dendritic/axonal-related functions (T(28)=-4.5612, p=9.197e-05; dendritic/axonal vs. non-dendritic/axonal: T(50.082)=2.3385, p=0.02339, **Suppl. Figure 11**). Based on this analysis, we conclude that synaptic atrophy could be partially driving age-related effects on neuronal cell type indices in the human prefrontal cortex dataset but are unlikely to fully explain the relationship.

Next, we decided to make the process of differentiating between altered cell type-specific functions and relative cell type balance more efficient. We used our cell type specific gene lists to construct gene sets in a file format (.gmt) compatible with the popular tool Gene Set Enrichment Analysis (39,40). Then, for the results from each subject variable within the Pritzker dataset, we compared the enrichment of the effects within gene sets defined by brain cell type to the enrichment seen within gene sets for other functional categories. In general, we found that gene sets for brain cell types tended to be the top result (most extreme normalized enrichment score, NES) for each of the subject variables that showed a strong relationship with cell type in our previous analyses (Agonal Factor vs. “Neuron_All_Cahoy_JNeuro_2008”: NES=-2.46, p=0.00098, q=0.012, Brain pH vs. “Astrocyte_All_Cahoy_JNeuro_2008”: NES=-2.48, p=0.0011, q=0.014, MDD vs. “Astrocyte_All_Cahoy_JNeuro_2008”: NES=-2.60, p=0.0010, q=0.017, PMI vs. “GO_OLIGODENDROCYTE_DIFFERENTIATION”: NES=-2.42, p=0.00078, q=0.027; **Suppl. Table 6**). Similarly, the relationship between the effects of age and neuron-specific gene expression was ranked #4, following the gene sets “GO_SYNAPTIC_SIGNALING”, “REACTOME_TRANSMISSION_ACROSS_CHEMICAL_SYNAPSES”, “REACTOME_OPIOID_SIGNALLING”, but each of them was assigned a similar p-value (p=0.001) and adjusted p-value (q=0.036). We conclude that it is important to consider cell type-specific expression during the analysis of macro-dissected brain microarray data above and beyond the consideration of specific functional pathways, and have submitted our .gmt files to the Broad Institute for addition to their curated gene sets in MSigDB to promote this form of analysis.

#### 3.11 Including Cell Content Predictions in the Analysis of Microarray Data Improves Model Fit and Enhances the Detection of Diagnosis-Related Genes in Some Datasets

Over the years, many researchers have been concerned that transcriptomic analyses of neuropsychiatric illness often produce non-replicable or contradictory results and, perhaps more disturbingly, are typically unable to replicate well-documented effects detected by other methods. We posited that this lack of sensitivity and replicability might be partially due to cell type variability in the samples, especially since such a large percentage of the principal components of variation in our samples were explained by neuron to glia ratio. Within the Pritzker dataset, we were particularly interested in controlling for cell type variability, because dissection may have differed between technical batches that were unevenly distributed across diagnosis categories (**Figure 5 A**). There was a similarly uneven distribution of dissection methods across diagnosis categories within the large CMC RNA-Seq dataset. In this dataset, the majority of the bipolar samples (75%) were collected by a brain bank that performed gray matter only dissections (PITT), whereas the control and schizophrenia samples were more evenly distributed across all three institutions (34).

**Figure 5.**
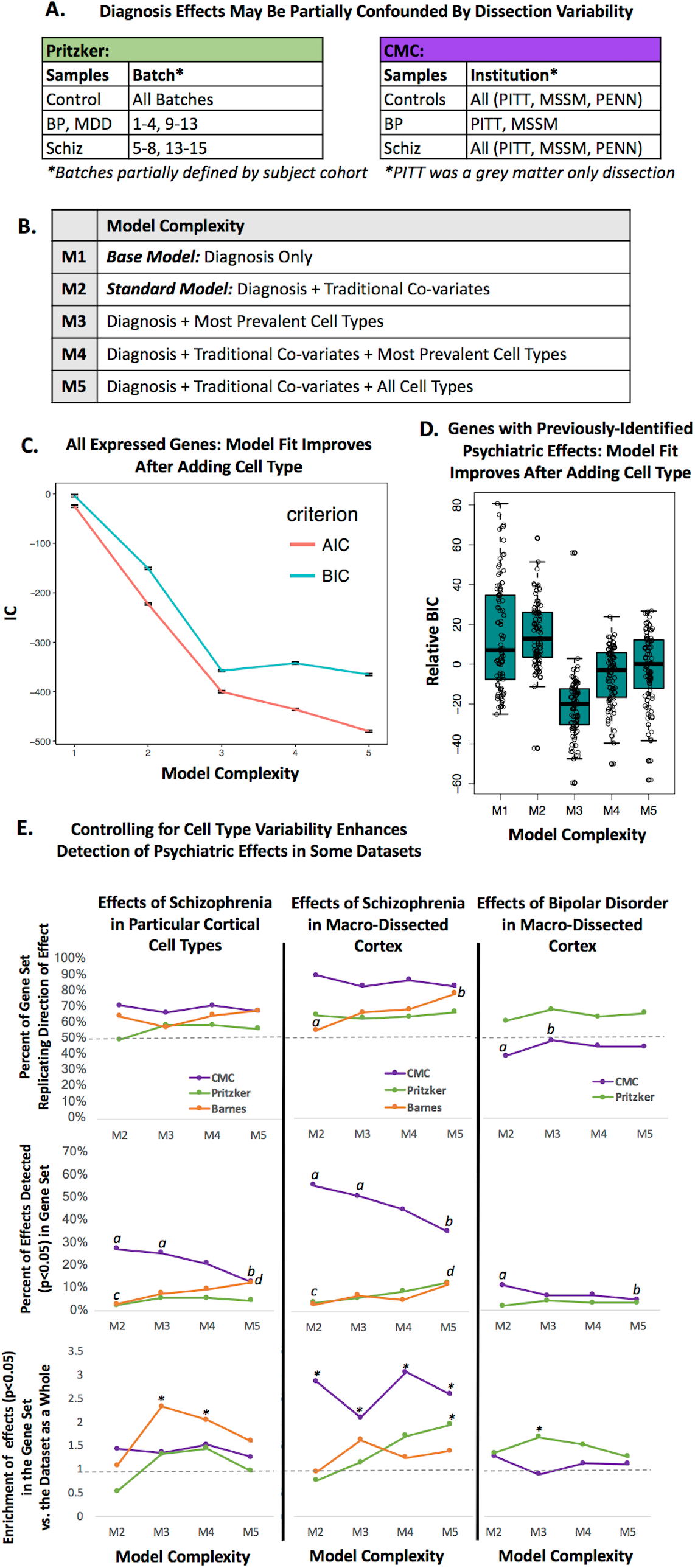
Including Cell Content Predictions in the Analysis of Microarray Data Improves Model Fit and Enhances the Detection of Previously-Identified Diagnosis-Related Genes in Some Datasets. **A.** Diagnosis effects were likely to be partially confounded by dissection variability within the Pritzker and CMC datasets. **B:** We examined a series of differential expression models of increasing complexity, including a base model (M1), a standard model (M2), and three models that included cell type co-variates (M3-M5). **C-D.** Model fit improved with the addition of cell type (M1/M2 vs. M3-M5) when examining either **C**. all expressed genes in the dataset (example from CMC: points= AVE +/-SE). **D.** genes with previously-documented relationships with psychiatric illness in particular cell types (example from Pritzker: BIC values for all models for each gene were centered prior to analysis. Boxes represent the median and interquartile range of the data). **E.** Evaluating the replication of previously-observed psychiatric effects (**Suppl. Figure 12**) in three datasets (Pritzker, CMC, and Barnes) using a standard differential expression model (M2) vs. models that include cell type co-variates (M3-5). Letters (a vs. b, c vs. d) denote significant model comparisons (Fisher’s exact test: p<0.05). Top graphs: The percentage of genes (y-axis: 0-1) replicating the direction of previously-documented psychiatric effects on cortical gene expression sometimes increases with the addition of cell type to the model (p<0.05: Barnes (effects of Schiz): M2 vs. M5, CMC (effects of Bipolar Disorder): M2 vs. M3). Middle graphs: The detection of previously-identified psychiatric effects on gene expression (p<0.05 & replicated direction of effect) increases with the addition of cell type to the model in some datasets (p<0.05, Barnes: M2 vs. M5, Pritzker: M2 vs. M5) but decreases in others (p<0.05, CMC: M2 vs. M5, M3 vs. M5). Bottom graphs: In some datasets we see an enrichment of psychiatric effects (*p<0.05) in previously-identified psychiatric gene sets only after controlling for cell type (Barnes: M3, M4, Pritzker: M5, M3). For the CMC dataset, we see an enrichment using all models (*p<0.05).

We hypothesized that controlling for cell type while performing differential expression analyses in these datasets would improve our ability to detect previously-documented psychiatric effects on gene expression, especially psychiatric effects that were previously-identified within specific cell types, since these effects should not be mediated by psychiatric changes in overall cell type balance. To test the hypothesis, we first compiled a list of 130 strong, previously-documented relationships between Schizophrenia or Psychosis and gene expression in particular cell types in the human cortex, as detected by *in situ* hybridization or immunocytochemistry ((70–75) reviewed further in (19)) or by single-cell type laser capture microscopy (**Suppl. Figure 12, Suppl. Table 7** (1,76,77)).

As a comparison, we also considered lists of transcripts strongly-associated with Schizophrenia (78) and Bipolar Disorder (79) in meta-analyses of microarray data derived from human frontal cortical tissue (**Suppl. Figure 12**, **Suppl. Table 7**). The effects of psychiatric illness on the expression of these transcripts could be mediated by either psychiatric effects on cell type balance or by effects within individual cells. Therefore, controlling for cell type balance while performing differential expression analyses could detract from the detection of some psychiatric effects, but perhaps also enhance the detection of other psychiatric effects by controlling for large, confounding sources of noise (*e.g.*, dissection variability).

Next, we examined our ability to detect these previously-documented psychiatric effects using regression models of increasing complexity (**Figure 5 B**), including a standard model controlling for traditional co-variates (Model 2) and models controlling for cell type co-variates (Models 3-5).

We found that including predictions of cell type balance in our models assessing the effect of diagnosis on gene expression dramatically improved model fit as assessed by Akaike’s Information Criterion (AIC) or Bayesian Information Criterion (BIC). These improvements were largest with the addition of the five most prevalent cell types to the model (M3, M4); the addition of less common cell types produced smaller gains (M5). These improvements were clear whether we considered the average model fit for all expressed genes (*e.g.*, **Figure 5 C**) or just genes with previously-identified psychiatric effects (*e.g.*, **Figure 5 D**).

However, models that included cell type were not necessarily superior at replicating previously-observed psychiatric effects on gene expression, even when examining psychiatric effects that were likely to be independent of changes in cell type balance. For each model, we quantified the percentage of genes replicating the previously-observed direction of effect in relationship to psychiatric illness, as well as the percentage of genes that replicated the effect using a common threshold for detection (p<0.05). Finally, we also looked at the enrichment of psychiatric effects (p<0.05) in each of the previously-documented psychiatric gene sets in comparison to the other genes in our datasets (genes universally represented in all three datasets-Pritzker, CMC, Barnes).

In general, we found that the two datasets that had the most variability in gene expression related to cell type (Pritzker, Barnes) were more likely to replicate previously-documented psychiatric effects on gene expression when the differential expression model included cell type covariates. For example, in the Barnes dataset, adding cell type co-variates to the model increased our ability to detect effects of Schizophrenia that had been previously documented within particular cell types or macro-dissected tissue (**Figure 5E**, Fisher’s exact test: M2 vs. M5, *p*<0.05 in both gene sets) and revealed an enrichment of Schizophrenia effects in genes with previously-documented psychiatric effects in particular cell types (Fisher’s exact test p<0.05: M3 & M4). In the Pritzker dataset, adding cell type co-variates to the model increased our ability to detect previously-documented effects of Schizophrenia in macrodissected tissue (M2 vs. M5: p<0.05) and revealed a significant enrichment of Schizophrenia and Bipolar effects in genes with previously-documented psychiatric effects in macro-dissected tissue (Fisher’s exact test p<0.05: Schizophrenia: M5, Bipolar: M3). This mirrored the results of another analysis that we had conducted suggesting that controlling for cell type increased the overlap between the top diagnosis results in the Pritzker dataset and previous findings in the literature as a whole (***Suppl. Section 7.3.4***).

In the large CMC RNA-Seq dataset, the rate of replication of previously-documented effects of Schizophrenia was already quite high using a standard differential expression model containing traditional co-variates (M2). Using a standard model, we could detect 27% of the previously-documented effects in cortical cell types and 55% of the previously-documented effects in macro-dissected tissue (with a replicated direction of effect and p<0.05). However, in contrast to what we had observed in the Pritzker and Barnes datasets, controlling for cell type *diminished* the ability to detect effects of Schizophrenia that had been previously-observed within particular cell types or macrodissected tissue in a manner that scaled with the number of co-variates included in the model (M2 or M3 vs. M5: p<0.05 for both gene sets), despite improvements in model fit parameters and a lack of significant relationship between Schizophrenia and any of the prevalent cell types. Including cell type co-variates in the model did not improve our ability to observe an enrichment of Schizophrenia effects in genes with previously-documented psychiatric effects in macro-dissected tissue (all models showed enrichment, M2-M5: Fisher’s exact test p<0.05). Controlling for cell type slightly improved the replication of the direction of previously-documented Bipolar Disorder effects (Fisher’s exact test: M2 vs. M3: p<0.05) in a manner that would seem appropriate due to the highly uneven distribution of bipolar samples across institutions and dissection methods, but even after this improvement the rate of replication was still no better than chance (48%), and, counterintuitively, the ability to successfully detect those effects still diminished in a manner that seemed to scale with the number of co-variates included in the model (Fisher’s exact test: M2 vs. M5, p<0.05). In a preliminary analysis of the two smaller human microarray datasets that were derived from gray-matter only dissections (GSE53987, GSE21138), the addition of cell type co-variates to differential expression models clearly diminished both the percentage of genes replicating the previously-documented direction of effect of Schizophrenia in particular cell types (Fisher’s exact test: GSE21138: M2 vs. M4 or M5: p<0.05, GSE53987: M2 vs. M4 or M5: p<0.05) and the ability to successfully detect previously-documented effects (Fisher’s exact test: GSE21138: M2 vs. M4 or M5: p<0.05).

#### General Discussion

We found that including cell type indices as co-variates while running differential expression analyses helped improve our ability to detect previously-documented relationships between psychiatric illness and gene expression in human cortical datasets that were particularly affected by variability in cell type balance. This improvement was not seen in datasets that were less affected by variability in cell type balance, despite improvements in model fit and a lack of strong multicollinearity between diagnosis and the cell type indices. This finding was initially surprising to us, but upon further consideration makes sense, as the cell type indices are multi-parameter gene expression variables.

Therefore, there is increased risk of overfitting when modeling the data for any particular gene. We conclude that the addition of cell type covariates to differential expression models is only recommended when there is a particularly large amount of variability in the dataset associated with cell type balance, or when there is strong reason to believe that technical variation associated with cell type (such as dissection) may be highly confounding in the result. We strongly recommend that model selection while conducting differential expression analyses should be considered carefully, and evaluated not only in terms of fit parameters but also validity and interpretability.

Regarding the importance of model selection for interpretability, it is worth noting that an important difference between our final analysis methods and those used by some previous researchers (*e.g.*, 10–12) was the lack of cell type interaction terms included in our models (*e.g.*, Diagnosis*Astrocyte Index). Theoretically, the addition of cell type interaction terms should allow the researcher to statistically interrogate cell-type differentiated diagnosis effects because samples that contain more of a particular cell type should exhibit more of that cell type’s respective diagnosis effect. Versions of this form of analysis have been successful in other investigations (e.g., (11,12,80)) but we were not able to validate the method using a variety of model specifications and our database of previously-documented relationships with diagnosis in prefrontal cell types. Upon consideration, we realized that these negative results were difficult to interpret because significant diagnosis*cell type interactions should only become evident if the effect of diagnosis in a particular cell type is different from what is occurring in all cell types on average. For genes with expression that is reasonably specific to a particular cell type (*e.g.*, GAD1, PVALB), the overall average diagnosis effect may already largely reflect the effect within that cell type and the respective interaction term will not be significantly different, even though the disease effect is clearly tracking the balance of that cell population. In the end, we decided that the addition of interaction terms to our models was not demonstrably worth the associated decrease in overall model fit and statistical power. For public use we have released the full differential expression results for each dataset analyzed using the different models discussed above (**Suppl. Table 8**-**Suppl. Table 12**).

## 4. Conclusion and Future Directions

In this manuscript, we have demonstrated that the statistical cell type index is a relatively simple manner of interrogating cell-type specific expression in transcriptomic datasets from macro-dissected human brain tissue. We find that statistical estimations of cell type balance almost fully account for the top principal components of variation in microarray data derived from macro-dissected brain tissue samples, far surpassing the effects of traditional subject variables (post-mortem interval, hypoxia, age, gender). Indeed, our results suggest that many variables of medical interest are themselves accompanied by strong changes in cell type specific gene expression in naturally-observed human brains. We find that within both chronic (age, sex, diagnosis) and acute conditions (agonal, PMI, pH) there may be substantial changes in the relative representation of different cell types. Thus, accounting for demography at the cellular population level can be as important for the interpretation of microarray data as cell-level functional regulation. This form of data deconvolution was useful for identifying the subtler effects of psychiatric illness within our samples, divulging the decrease in astrocytes that is known to occur in Major Depressive Disorder and the decrease in red blood cell content in the frontal cortex in Schizophrenia, resembling known fMRI hypofrontality. This form of data deconvolution may also aid in the detection of psychiatric effects while conducting differential expression analyses in datasets that have highly-variable cell content.

These results touch upon the fundamental question as to whether organ-level function responds to challenge by changing the biological states of individual cells or the life and death of different cell populations. To reach such a sweeping perspective in human brain tissue using classic cell biology methods would require epic efforts in labeling, cell sorting, and counting. We have demonstrated that scientists can approximate this vantage point using an elegant, supervised signal decomposition exploiting increasingly available genomic data. However, it should be noted that, similar to other forms of functional annotation, cell type indices are best treated as a hypothesis-generation tool instead of a final conclusion regarding tissue cell content. We have demonstrated the utility of cell type indices for detecting large-scale alterations in cell content in relationship with known subject variables in post-mortem tissue. We have not tested the sensitivity of the technique for detecting smaller effects or the validity under all circumstances or non-cortical tissue types. Likewise, while using this technique it is impossible to distinguish between alterations in cell type balance and cell-type specific transcriptional activity: when a sample shows a higher value of a particular cell type index, it could have a larger number of such cells, or each cell could have produced more of its unique group of transcripts, via a larger cell body, slower mRNA degradation, or an overall change in transcription rate. In this regard, the index that we calculate does not have a specific interpretation; rather it is a holistic property of the cell populations, the “neuron-ness” or “microglia-ness” of the sample. Such an abstract index represents the ecological shifts inferred from the pooled transcriptome. That said, our cell type indices do have real biological meaning - they can be interpreted in a known system of cell type taxonomy. When single-cell genomic data uncovers new cell types (e.g., the Allen Brain Atlas cellular taxonomy initiative (81)) or meta-analyses refine the list of genes defined that have cell-type specific expression (e.g., (82)), our indices will surely evolve with these new classification frameworks, but the power of the approach will remain, in that we can disentangle the intrinsic changes of individual genes from the population-level shifts of major cell types.

Our work drives home the fact that any comprehensive theory of psychiatric illness needs to account for the dichotomy between the health of individual cells and that of their ecosystem. We found that the functional changes accompanying psychiatric illness in the cortex occurred both at the level of cell population shifts (decreased astrocytic presence and red blood cell count) and at the level of intrinsic gene regulation not explained by population shifts. A similar conclusion regarding the importance of cell type balance in association with psychiatric illness was recently drawn by our collaborators (e.g.,(83)) using a similar technique to analyze RNA-Seq data from the anterior cingulate cortex. In the future, we plan to use our technique to re-analyze many other large transcriptomic datasets with the hope of gaining better insight into psychiatric disease. This application of our technique seems particularly important in light of recent evidence linking disrupted neuroimmunity (84) and neuroglia (e.g., (48,57,85)) to psychiatric illness, as well as growing evidence that growth factors with cell type specific effects play an important role in depressive illness and emotional regulation (for a review see (23,86)).

In conclusion, we have found this method to be a valuable addition to traditional functional ontology tools as a manner of improving the interpretation of transcriptomic results. For the benefit of other researchers, we have made our database of brain cell type specific genes (**Suppl. Table 1**, https://sites.google.com/a/umich.edu/megan-hastings-hagenauer/home/cell-type-analysis) and code for conducting cell type analyses publicly available in the form of a downloadable R package (https://github.com/hagenaue/BrainInABlender) and we are happy to assist researchers in their usage for pursuing better insight into psychiatric illness and neurological disease.

## 5. Acknowledgements

We thank all members of the Pritzker Consortium (especially the University of California, Irvine Brain Bank staff), Drs. Adriana Medina and David Krolewski for brain dissections and methodological input, and Dr. Simon Evans, Sharon Burke and Mary Hoverstein for their involvement in the initial mRNA extraction and microarrays. Grace Hsienyuan Chang, Jennifer Fitzpatrick, LeAnn Fitzpatrick, Jim Stewart, Tom Dixon, Doug Smith, Andy Lin, and Manhong Dai were invaluable for maintaining our databases of clinical information and biological specimens. Drs. Elyse Aurbach, Katherine Prater, Kathryn Hilde, Fan Meng, Lilah Toker, Mark Reimers, and Angela O’Connor provided insightful advice and feedback regarding the methodology or manuscript. Our undergraduate research assistants Isabelle Birt, Alek Pankonin, and Daniela Romero Vargas helped compile the Allen Brain Atlas data, annotate and upload code, create the BrainInABlender R package, and provide editorial assistance. Finally, we would like to thank our reviewers, whose insightful feedback helped inspire several particularly useful analyses, leading us to a stronger set of conclusions.

## Footnotes

* As of 9-14-2017

